# Mapping the Immunodominance Landscape of SARS-CoV-2 Spike Protein for the Design of Vaccines against COVID-19

**DOI:** 10.1101/2020.04.23.056853

**Authors:** Bao-zhong Zhang, Ye-fan Hu, Lin-lei Chen, Yi-gang Tong, Jing-chu Hu, Jian-piao Cai, Kwok-Hung Chan, Ying Dou, Jian Deng, Hua-rui Gong, Chaiyaporn Kuwentrai, Wenjun Li, Xiao-lei Wang, Hin Chu, Cai-hui Su, Ivan Fan-Ngai Hung, Thomas Chung Cheung Yau, Kelvin Kai-Wang To, Kwok Yung Yuen, Jian-Dong Huang

## Abstract

The ongoing coronavirus disease 2019 (COVID-19) pandemic is a serious threat to global public health, and imposes severe burdens on the entire human society. The severe acute respiratory syndrome (SARS) coronavirus-2 (SARS-CoV-2) can cause severe respiratory illness and death. Currently, there are no specific antiviral drugs that can treat COVID-19. Several vaccines against SARS-CoV-2 are being actively developed by research groups around the world. The surface S (spike) protein and the highly expressed internal N (nucleocapsid) protein of SARS-CoV-2 are widely considered as promising candidates for vaccines. In order to guide the design of an effective vaccine, we need experimental data on these potential epitope candidates. In this study, we mapped the immunodominant (ID) sites of S protein using sera samples collected from recently discharged COVID-19 patients. The SARS-CoV-2 S protein-specific antibody levels in the sera of recovered COVID-19 patients were strongly correlated with the neutralising antibody titres. We used epitope mapping to determine the landscape of ID sites of S protein, which identified nine linearized B cell ID sites. Four out of the nine ID sites were found in the receptor-binding domain (RBD). Further analysis showed that these ID sites are potential high-affinity SARS-CoV-2 antibody binding sites. Peptides containing two out of the nine sites were tested as vaccine candidates against SARS-CoV-2 in a mouse model. We detected epitope-specific antibodies and SARS-CoV-2-neutralising activity in the immunised mice. This study for the first time provides human serological data for the design of vaccines against COVID-19.

## Introduction

The coronavirus disease 2019 (COVID-19), a novel infectious disease caused by severe acute respiratory syndrome coronavirus 2, SARS-CoV-2, first emerged in December 2019 and has since become a worldwide pandemic. COVID-19 infections have so far led to over 2 million cases and more than 0.13 million deaths across more than 200 countries and geographical regions around the world. The World Health Organization has launched a worldwide clinical trial called SOLIDARITY to test antiviral treatment options for the novel coronavirus, but as of April 15, 2020, there are no specific drugs for treating COVID-19^1^. Many research teams worldwide are racing to develop vaccines against SARS-CoV-2 to combat this novel coronavirus pandemic.

Similar to other coronaviruses, SARS-CoV-2 is an enveloped positive-stranded RNA virus with four major structural proteins: S (spike), E (envelope), M (membrane), and N (nucleocapsid) proteins^2-4^. The surface S protein is the key that allows SARS-COV-to enter into cells^5^, as it plays a role in binding to the cellular receptor and membrane fusion^5,6^. The SARS-CoV-2 shares 75.96% homology with the 2003 SARS-CoV^3^, and the structure and functional domains of the SARS-CoV-2 S protein have already been identified^7^. The receptor-binding domain (RBD) within the S1 domain binds to angiotensin-converting enzyme 2 (ACE2) to facilitate the entry of SARS-CoV-2 into host cells^6,7^.

As of April 8, 2020, there are at least 115 vaccine candidates in active development worldwide^8^. The SARS-CoV-2 S (spike) glycoprotein is the immunogen that is the focus of the majority of the vaccine research. In the absence of *in vivo* and *in vitro* data, researchers have been using *in silico* data based on the IEDB database or other online epitope prediction algorithms^9-14^. However, the *in silico* data is less likely to be useful for vaccine development, because these bioinformatics tools were not optimised for vaccine design. Furthermore, the antigenic properties of S proteins remain elusive, and therefore experimental immunogenic information is urgently needed.

Although a putative epitope that binds to antibodies against SARS-CoV-2 has been reported^15^, more epitopes with high affinity need to be identified. Information on these epitopes would be useful, particularly in regard to human immunological bias and immunodominance (ID), which might restrict the immune response. Immunodominance is the phenomenon of immunogenic variation among distinct immunogens or antigenic sites on the same immunogen, which has been demonstrated in both CD8^+^ T cells and B cells^16^. This phenomenon has restricted the development of effective vaccines for influenza A and other highly variable viruses^17-19^. All the methods used to overcome or manipulate these viral immunity restrictions rely on the landscape of immunodominance. It is therefore crucial to map the immunogenicity landscape of the potential epitopes of the S protein to accelerate the development of vaccines.

An ideal vaccine against SARS-CoV-2 should induce highly potent neutralising antibodies but should not induce any disease-enhancing antibodies. Previous studies that used the full-length S protein of SARS-CoV to immunise animals resulted in adverse effect after the animals were challenged with SARS-CoV^20^. Jiang et al., identified five linear immunodominant (ID) sites in the S protein that did not induce neutralising antibodies. On the contrary, the RBD was found to contain the major neutralising epitopes in the S protein, but did not have any ID sites^21^.

In this study, we aimed to map the landscape of ID sites within the S protein of SARS-CoV-2 and compare these results with those of its counterpart in SARS-CoV. Unlike SARS-CoV, we found the RBD of SARS-CoV-2 had linear ID sites in sera samples collected from recently discharged COVID-19 patients. Subsequent microneutralisation tests demonstrated that mice immunised with peptides containing the ID sites within RBD produced neutralising antibodies. Taken together, the RBD of SARS-CoV-2 has a different ID landscape to its counterpart in SARS-CoV. Furthermore, these data can provide guidance on the design of S protein-based SARS-CoV-2 vaccines that might avoid adverse reactions or disease-enhancing effects.

## Materials and Methods

### Serum specimens from COVID-19 patients

Serum samples were collected from 39 patients with COVID-19 (22 males and 17 females) between Jan 26, 2020, and March 18, 2020. Twenty-six patients who were discharged from hospital provided written informed consent under UW 13-265. The other 13 hospitalized patients were sampled under UW 13-372, and written informed consent was waived. Average sampling time from the onset date was 28.7 days (range, 14–44 days). The median age of patients was 59.7 years (range, 26-80 years). All patients were diagnosed with COVID-19 by PCR test, and two patients had a severe illness. The diagnostic criteria for SARS-CoV infection followed the clinical description of COVID-19^22^. The initial laboratory confirmation was performed on nasopharyngeal or sputum specimens at the Public Health Laboratory Centre of Hong Kong. The discharge criteria were clinically stable and negative nucleic acid test twice consecutively (sampling interval ≥ 24 hours). Six healthy donors were also sampled under UW 19-470. This study was approved by the Institutional Review Board of the University of Hong Kong/Hospital Authority Hong Kong West Cluster (UW 13-372, UW 13-265, and UW 19-470).

### Animal experiments

The SPF BALB/c mice were supplied by the Laboratory Animal Unit of the University of Hong Kong. All animal experiments were approved by the Committee on the Use of Live Animals in Teaching & Research, the University of Hong Kong (CULATR 5312-20). All mice were immunised by distinct vaccines on Day 0 and Day 14.

### Synthesis of peptides

All peptides were manufactured by GL Biochem (Shanghai) Ltd in the form of a dry powder. The peptides were generated using solid phase synthesis methods and the quality of the products was monitored by mass spectrometry. All peptides were dissolved in pH 7.4 PBS buffer or 8 M pH 7.0 urea Na2HPO4/NaH2PO4 buffer. Peptides conjugated to keyhole limpet haemocyanin (KLH) carrier proteins were also purchased from GL Biochem (Shanghai) Ltd. These peptide-conjugated proteins were dissolved in a pH 7.4 PBS buffer.

### Antibody detection using ELISA

Epitope-specific antibodies were detected by enzyme-linked immunosorbent assay (ELISA). Briefly, all peptides at a final concentration of 0.5 μg/mL in 50 mM pH 9.6 Na2CO3/NaHCO3 buffer were coated on ELISA plates (Nunc, Roskilde, Denmark) and incubated overnight at 4°C. Plates were blocked with TBS −5% (w/v) non-fat milk for 3 h at 37°C and washed four times in 0.05% Tween in TBST. Diluted patient or mice sera were added into the wells and incubated for 1 h at 37°C. Plates were washed six times in TBS-0.05% Tween and incubated with HRP-conjugated goat anti-mouse IgG for 1 h at 37°C. The colour was developed using Trimethyl Borane (TMB) solution (Sigma) and absorbance was measured at 450 nm using an ELISA reader. Samples from non-immunised mice or healthy volunteers were used as the controls. The cut-off lines were based on the mean value plus three times the standard deviation.

### Identification of T cell epitopes using ELISpot

The T cell responses were detected by enzyme-linked immunosorbent spot (ELISpot) kits (Dakewe Biotech Co., Ltd) following the manufacturer’s protocol. Briefly, splenocytes harvested from sacrificed mice were washed and immediately transferred to anti-IFN-γ antibody pre-coated filter plates. For stimulation, splenocytes were co-incubated with distinct epitopes overnight at 37°C. All samples were assayed with positive controls (Phorbol myristate acetate, PMA and Ionomycin) and cells from a reference donor. All images in different wells were captured using a CTL ImmunoSpot ELISpot Analyzer and processed using the ImmunoCapture software (Cellular Technology Ltd., USA).

### Microneutralisation tests

Serial two-fold dilutions of heat-inactivated sera (treated at 56°C for 30 minutes) were prepared from a starting dilution of 1:10. The serum dilutions were mixed with equal volumes of 100 TCID50 of SARS-CoV-2 as indicated. After 1 h of incubation at 37°C, 35 µL of the virus–serum mixture was added to Vero-E6 cell monolayer for SARS-CoV-2 infection in 96-well microtitre plates in quadruplicate. After 1 h of adsorption, an additional 150 µL of culture medium was added to each well and incubated for 3 days at 37°C in 5% CO2 in a humidified incubator. A virus back-titration was performed without immune serum to assess the input virus dose. The CPE was read at days post infection. The highest serum dilution that completely protected cells from CPE in half the wells was estimated using the Reed-Muench method and was taken as the neutralising antibody titre. Positive and negative control sera were included to validate the assay.

## Results

### Immune responses and antiviral effects in serum from COVID-19 patients

All serum samples from COVID-19 patients tested positive for SARS-CoV-2 by ELISA assay using plates coated by SARS-CoV-2 lysates (Figure 1A). The profiles of IgG, and IgM, and IgA against S (spike) and N (nucleocapsid) proteins showed that not all patients had elevated antibody titres compared to healthy donors (Figure 1B-G). Generally, high IgG titres against both S and N proteins were present in the sera, samples, and relatively high IgM and IgA responses were also observed in the majority of patients.

**Figure 1.**
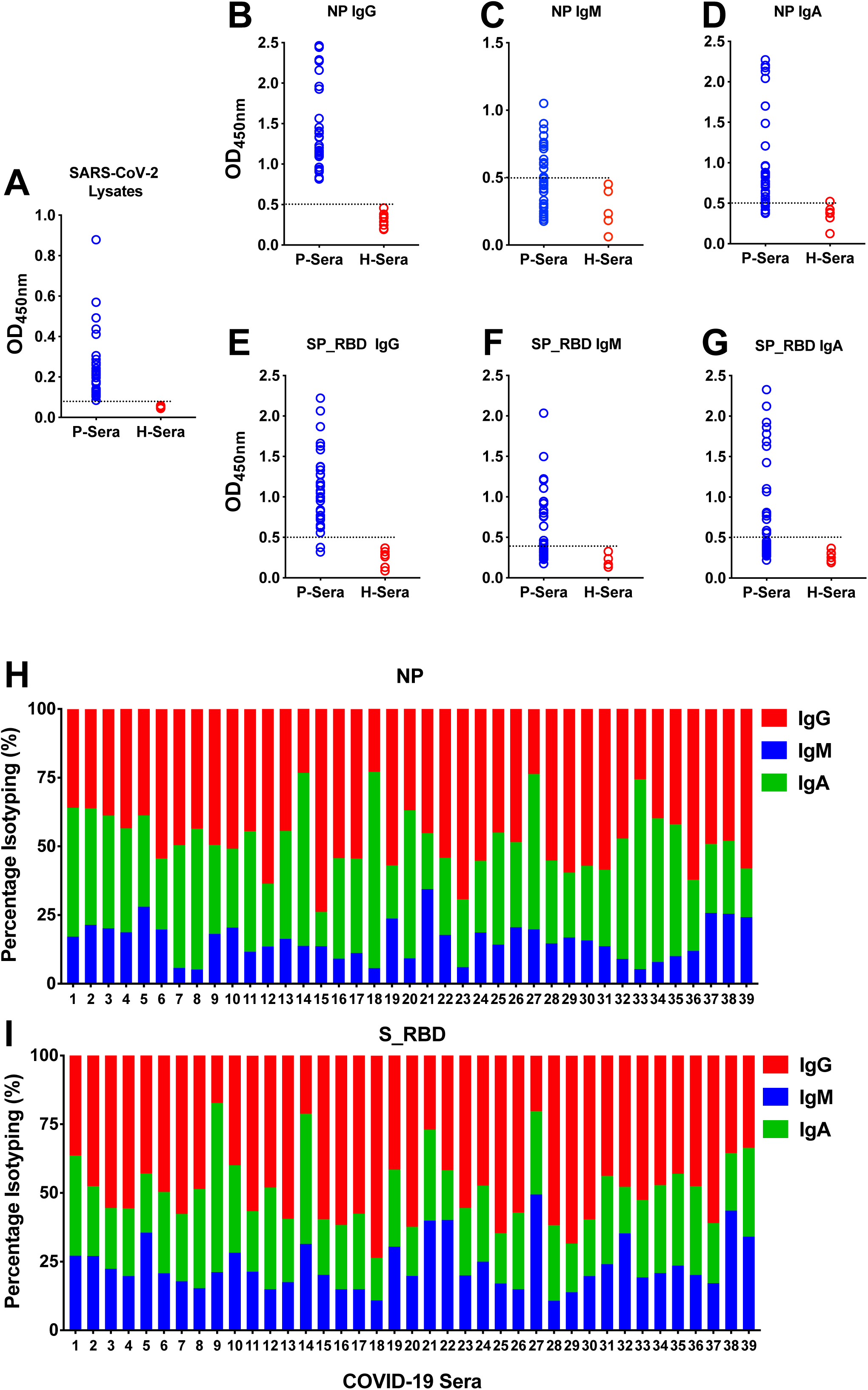
Detection of antibodies specific for SARS-CoV-2 proteins in early convalescent sera from COVID-19 patients by ELISA. (A) Total proteins from SARS-CoV-2 lysates were used as the coated antigen. (B-D) The recombinant N protein was used as the coated antigen. (E-G) The recombinant SP_RBD protein was used as the coated antigen. (H-I) Antibody isotyping of N and SP_RBD binding antibodies in early convalescent sera from COVID-19 patients. Sera from 39 COVID-19 patients, and 6 healthy blood donors were tested at a dilution of 1:100. The dashed lines represent cut-off values (the mean absorbance at 450 nm of sera from healthy blood donors plus three times the standard deviation).

Our previous work revealed that the correlations between microneutralisation (MN) activity and anti-NP or anti-RBD IgG titres were stronger than the correlation between MN activity and IgM titre^22^. To further investigate the correlations between MN activity and anti-NP or anti-RBD IgG, IgM, and IgA titres, we carefully analysed the sera from recovered patients. We found MN activity in the sera of 26 out of 39 COVID-19 patients. The MN titres were adjusted following previous criteria: MN titres less than 10 were re-designated a value of 5 and MN titres greater than 320 were re-designated a value of 640^22^. We identified a very strong correlation between anti-RBD IgG titres and MN activity in recovered patients (R^2^ = 0.8009). The correlations between anti-RBD IgM/IgA titres and MN activity were weaker than for IgG (R^2^ = 0.5130 and 0.5926, respectively) (Figure 2E, G, I). However, we found very poor correlations between anti-NP IgG/IgM/IgA titres and MN activity. These correlations dropped dramatically when MN activity titres were greater than 1:160 (Figure 2D, F, H). These results suggest that anti-RBD antibodies play an important role in the antiviral immune response.

**Figure 2.**
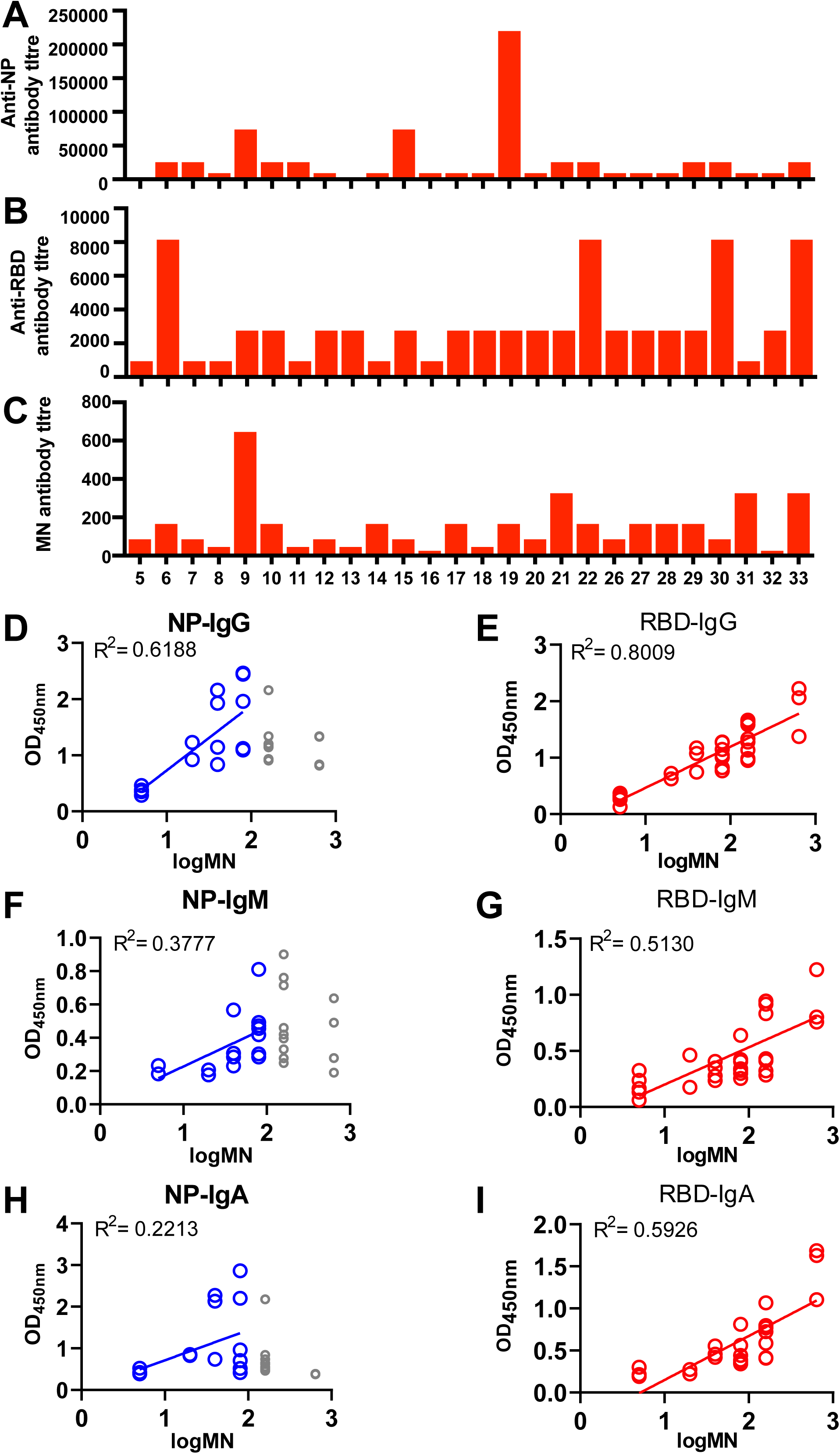
Correlations between S or N protein-specific antibody titres and microneutralisation antibody titres. (A-C) N protein or RBD fragment of S protein-specific IgG levels and microneutralisation (MN) assay results of recovered patients’ antibody titres. (D-E) Correlation between N protein or RBD fragment of S protein-specific IgG levels and microneutralisation antibody titres. (F-G) Correlation between N protein or RBD fragment of S protein-specific IgM levels and microneutralisation antibody titres. (H-I) Correlation between N protein or RBD fragment of S protein-specific IgA levels and microneutralisation antibody titres.

### The landscape of immunodominant sites on the S protein in COVID-19 patients

To identify the immunodominant (ID) sites on the SARS-CoV-2 S protein, we mapped the epitopes in 42 peptides spanning the entire extra-membrane domain (21-926) of the S protein with three gaps (106-160, 365-374, and 687-741). Each peptide was between 20 and 25 residues in length with a five-residue overlap. We measured ID sites in terms of the positive rate and the percentage of convalescent sera from COVID-19 patients having positive reactions to the epitope. Here, we used the mean response plus three times the standard deviation in healthy donors as the cut-off value to define positive reactions. The epitope mapping showed nine linear ID sites on the S protein located at 21-45(IDa), 221-245(IDb), 261-285(IDc), 330-349(IDd), 375-394(IDe), 450-469(IDf), 480-499(IDg), 522-646(IDh), and 902-926(IDi), respectively (Figure 3), with an average positive rate of ≥50% among all 39 patients. We found the SARS-CoV-2 RBD contained four ID sites, IDd, IDe, IDf, and IDg, whereas the SARS-CoV RBD has no ID sites. Considering the SARS-CoV-2 S protein shares 75.96% identity with the SARS-CoV S protein, we found five out of the nine fragments, IDc (79.17%), IDd (90%), IDe (90%), IDh (79.2%), and IDi (96%) were evolutionarily highly conserved in the SARS-CoV S protein. However, only three of the nine ID sites, IDa (63.64%), IDh (79.05%), and IDi (96%) were highly homologous (>50%) to SARS-CoV S protein ID sites. These results suggest that the conserved regions contribute to the immunogenicity of the S protein, whereas the majority of ID sites (6 out of 9) were less likely to be conserved. Interestingly, some epitopes induced personalized immune responses in specific patients. Patient 33 had an extremely strong immune response to epitope S (21-45) with a 40 times increase compared to the cut-off value, whereas the responses of patients 26 to 31 were not significantly different compared to those of healthy donors. A similar response was also observed for S (281-305), which is less likely to be an ID site. (Figure 3A) These results suggest that some unexpected ID sites of the S protein might lead to highly variable immune responses in patients if immunised with certain viral proteins.

**Figure 3.**
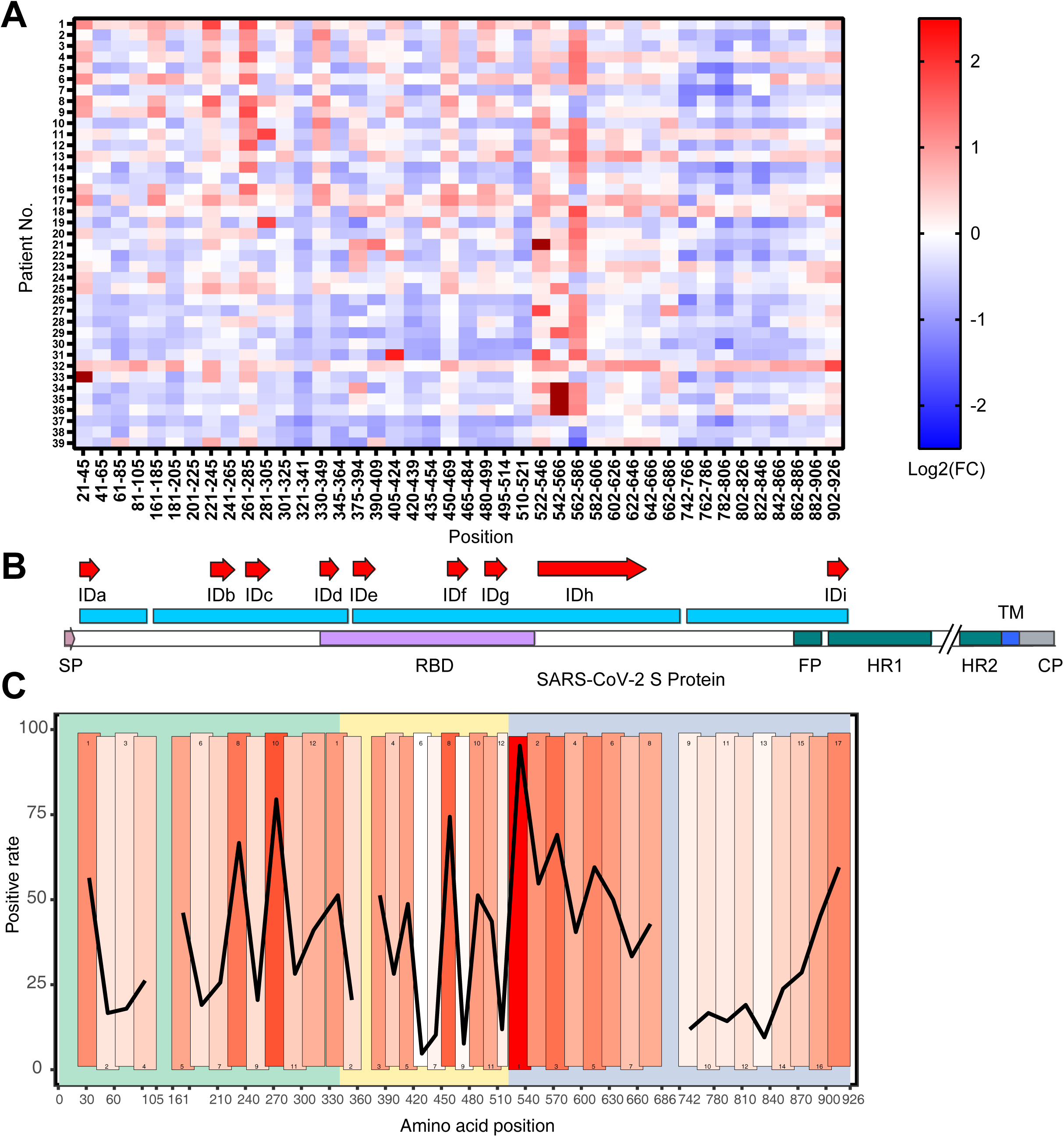
Epitope landscape of SARS-CoV-2 S proteins in convalescent sera from COVID-19 patients by ELISA. (A) The landscape of adjusted epitope-specific antibody levels in each patient. The ELISA results of absorbance at 450 nm were normalized to the aforementioned cut-off values. (B) Schematic representation of SARS-CoV-2 S protein and identified immunodominant sites. Here, only epitopes with positive rates greater than 50% are immunodominant. (C) Positive rates of distinct epitopes of SARS-CoV-2 S protein.

### Immunodominant sites can generate antiviral protection in a mouse model

To examine if specific ID sites of the S protein can generate immune responses and antiviral effects, we immunised mice with the RBD and two epitopes (S370-395 and S435-479) of the S protein, which were selected based on the results of B cell epitope prediction, toxicity prediction, and allergenicity prediction (Figure 4A, B). The S370-395 and S435-479 epitopes were validated in our epitope mapping assay in patient sera, which showed positive rates of 51.3% and 74.4%, respectively. The ELISA assay showed that mice immunised with the entire RBD, S370-395, and S435-479 generated high levels of specific antibodies (Figure 4C, D, E). In the MN test, mice immunised with S370-395, S435-479 or RBD showed viral neutralising titres of 1:26.7, 1:16.7, and 1:33.3, respectively (Figure 4F).

**Figure 4.**
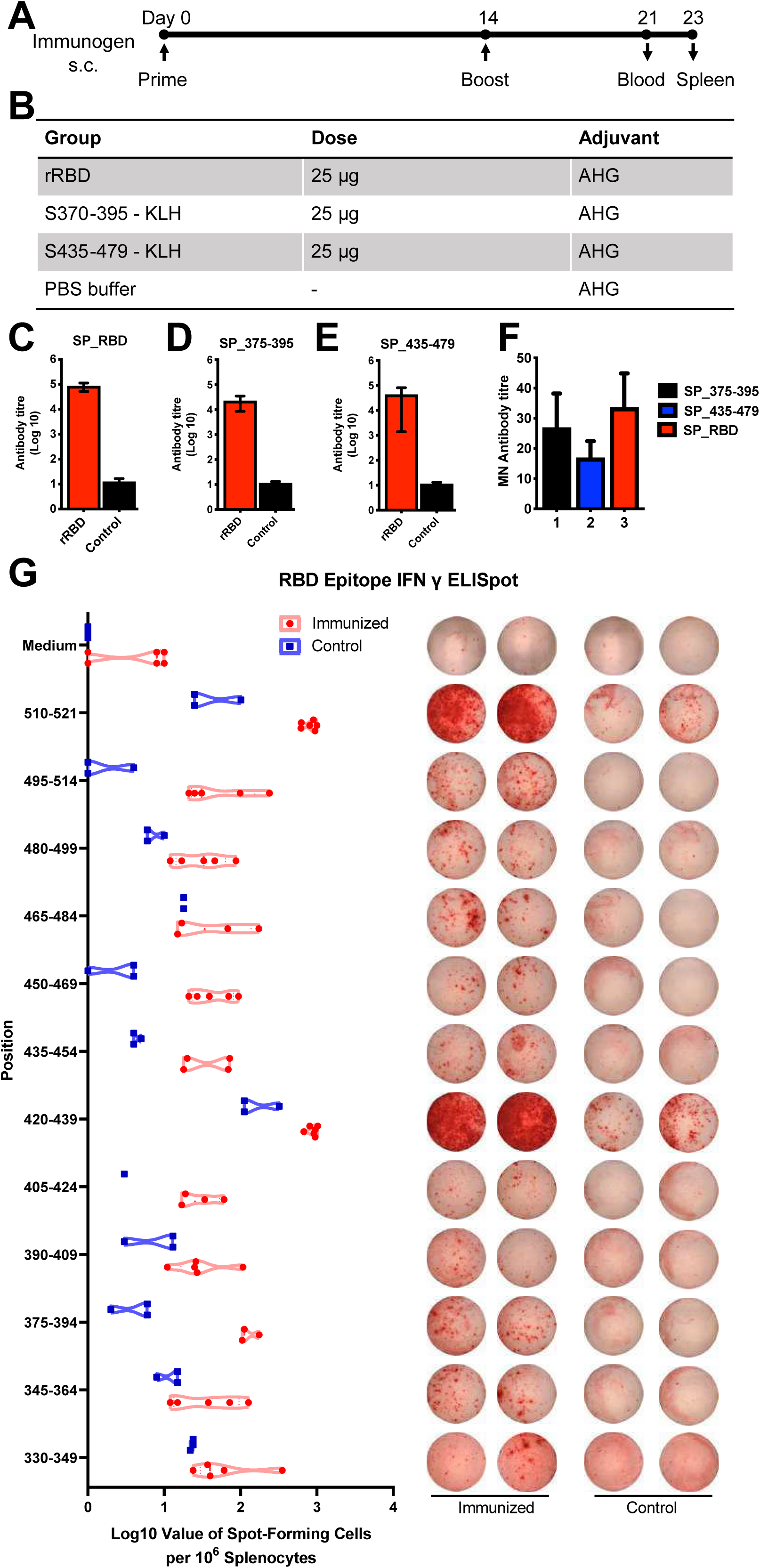
B Cell and T Cell responses to distinct epitopes in SARS-CoV-2 immunised mice. (A-B) Schematic representation of the mouse immunisation schedule. Balb/C mice (n = 5 per group) were immunised subcutaneously (s.c) with 25 µg of rRBD and KLH-conjugated peptides S370-395 and S435-479 mixed with aluminium hydroxide gel (AHG). (C-E) rRBD-specific antibody responses in immunised mice. rRBD-specific IgG antibody responses in mouse sera collected at 7 days after the second vaccination. (F) Virus microneutralisation antibody titres were measured against SARS-CoV-2 in classical BSL3. (G) Number of IFN-γ-secreting splenocytes in response to stimulation with the 12 RBD peptide pools of 20-mer peptides. SFU: spot-forming units

The landscape of T cell epitopes in the RBD fragment was profiled by ELISpot assay, which revealed a distinct pattern compared to the B cell responses. In the RBD fragment, three of the T cell epitopes, S405-469, S480-499, and S510-521, induced strong adaptive responses after immunisation. Among the epitopes, S370-395, S450-469 and S480-499 were identified as ID sites in human sera (Figure 4G). The S370-395 and S435-479 epitopes are more likely to be both T cell ID sites and B cell ID sites.

### Immunodominant sites might reveal potent neutralising sites of S protein

To validate the potential functions of ID sites, we highlighted the ID sites in the SARS-CoV-2 S protein structure model containing glycosylation sites^23^ (Figure 5A-C). Among ID sites with positive rates ranging from 50% to 60%, five sites were located on the head region of the S protein and two sites were on the stem of the S protein (Figure 5B). Among ID sites with positive rates over 60%, all five epitopes were located on the head of the S protein (Figure 5C). Although the S protein is a glycoprotein, glycosylation modification had limited effects on immunisation. Two glycosylation sites, N331 and N343, were observed in the RBD region, but these sites had no influence on the immune response. In sera from patients, non-glycosylated peptides were able to induce immune responses (Figure 3A, C and Figure 4G, S330-349).

**Figure 5.**
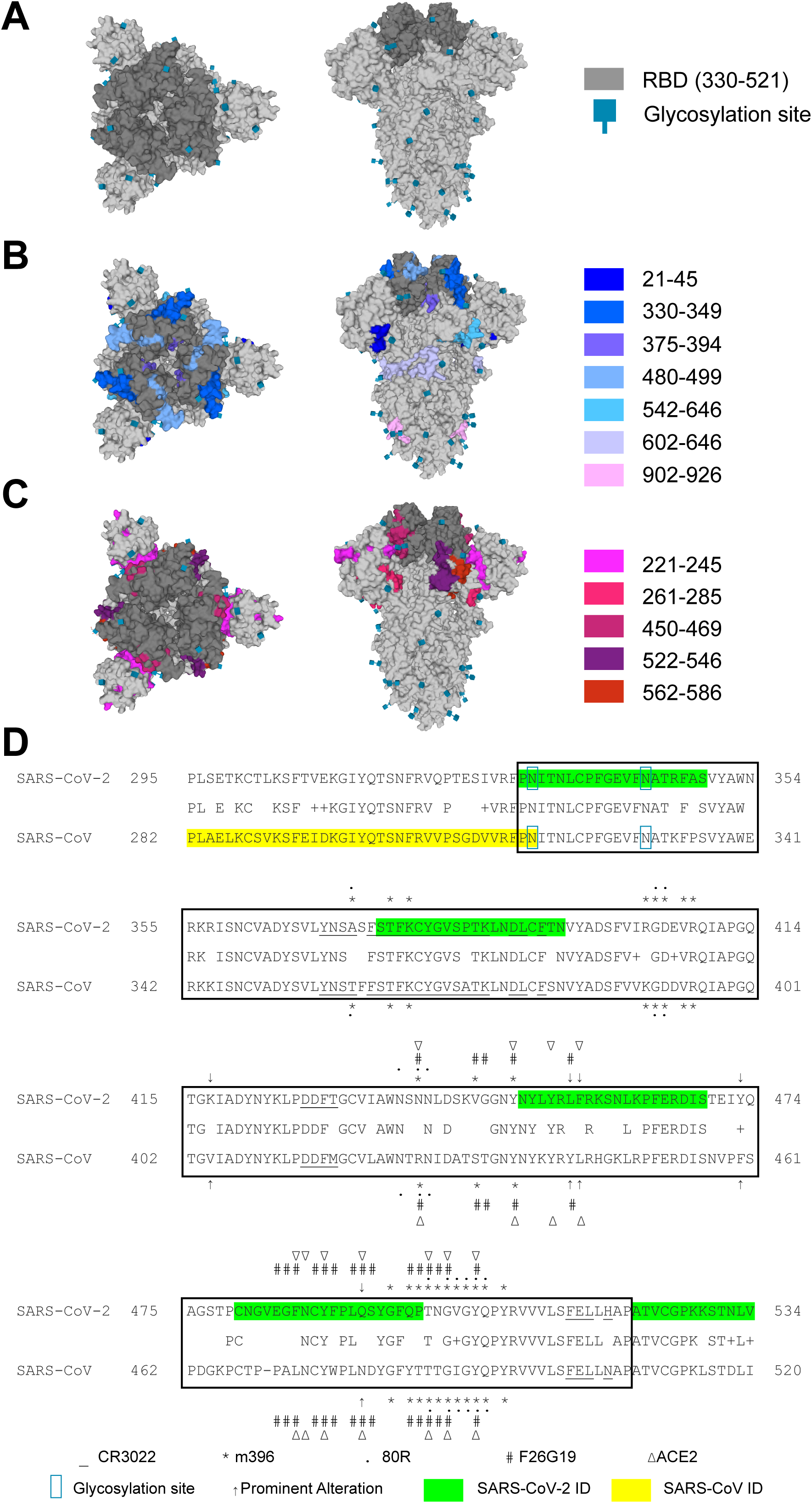
Structural representation of immunodominant and potential neutralising sites in SARS-CoV-2 RBD. (A) Top view and side view of the S (spike) protein in trimer form with labelled RBD region and glycosylation sites. (B) Top view and side view of the S (spike) protein with immunodominant sites with positive rates ranging from 50% to 60%. (C) Top view and side view of the S (spike) protein with immunodominant sites with positive rates greater than 60%. (D) Amino acid sequence alignment of SARS-CoV-2 and SARS-CoV RBD sequences. Related neutralising antibody binding sites, ACE2 binding sites, prominent alterations, and immunodominant sites of SARS-CoV-2 and SARS-CoV are shown.

The neutralising titre results (Figure 4F) suggest the ID sites have neutralising activities against SARS-CoV-2, although the ID sites in SARS-CoV were unlikely to be neutralising sites^20^. To further validate the ID sites, we compared the binding residues of the S protein to reported monoclonal antibodies with the ID sites of RBD^15,24,25^. Interestingly, some binding residues of SARS-CoV-2-specific neutralising antibodies were highly similar to the ID sites (Figure 5). For CR3022, a monoclonal antibody targeting a highly conserved cryptic epitope^15^, 14 out of 28 binding residues of SARS-CoV-2 RBD were located in ID sites. For F26G19, a mouse antibody, 11 out of 20 sites were located at ID sites, indicating there was likely a relatively high binding affinity to the RBD fragment^25^. However, for m396, an antibody with relatively low binding affinity to SARS-CoV-2 RBD, only 5 out of 22 binding residues were located at ID sites^24^. This was also the case for R80, another low binding affinity antibody, which had no matching binding residues^24^. These results suggest that SARS-CoV-2 ID sites are potent neutralising sites for high-affinity antibodies.

## Discussion

In this study, we profiled IgG/IgM/IgA levels against the S protein and N protein in the sera of COVID-19 patients (Figure 1). All convalescent sera from the COVID-19 patients contained specific antibodies against recombinant SARS-CoV-2 N protein, but not all sera had specific antibodies for the RBD fragment of the S protein. The relatively high immunogenicity of SARS-CoV-2 N protein during infection showed it has potential as an antigen for developing COVID-19 diagnostics (Figure 1). However, amounts of the different antibodies varied across patients. We found that IgM contributed 5%-34% of N protein-specific antibodies, whereas anti-RBD IgM contributed 10%-49% of RBD-specific antibodies. Patients with acute SARS-CoV-2 infection displayed highly diverse immune responses, and this diversity remained until convalescence. These diverse immune responses highlight safety concerns such as secondary infections that need to be considered.

We also analysed the correlation between S or N protein-specific antibody levels and SARS-CoV-2 neutralising titres. The Spike RBD-specific antibody level displayed a strong linear correlation with MN titres, but not with the N-specific antibody level (Figure 2). This observation suggests that RBD-specific antibodies in the sera of recovered patients might provide antiviral protection mainly through neutralising activity rather than non-neutralising antibodies against the N protein. This suggests that manipulating the RBD-induced immune responses might be more effective for developing COVID-19 vaccines.

This is the first reported mapping of the landscape of the ID sites in the S protein of SARS-CoV-2 using sera from COVID-19 patients (Figure 3). The personalized immune response pattern needs to be further investigated as unexpected and highly variable immune responses in some individuals might lead to adverse events or disease-enhancing responses to certain viral proteins used as vaccines. We further tested if the ID sites in the RBD fragment could be used as potential vaccines in mice. We found that epitopes/protein-specific antibody titres exceeded the microneutralisation titres by several orders of magnitude. Considering the relatively low neutralising antibody levels in the recovered patients in this study and in a previous report^26^, it is reasonable to expect significant differences between specific antibodies and neutralising antibodies. We found there was equivalent antiviral activity from immunisation with epitopes compared to immunisation with the entire RBD fragment (Figure 4). This provides evidence that epitope-based vaccines could offer comparable protection compared to subunit vaccines, but with the potentially better safety.

We also compared binding residues of previously identified SARS-CoV-2 neutralising antibodies with ID epitopes. The ID sites in the RBD were found to be potent neutralising sites for SARS-CoV-2 (Figure 5). Previous research suggests that it might be better to overcome the immunodominance of non-neutralising antigenic epitopes for SARS-CoV^20,27^, influenza virus A^16^, dengue fever virus^28^, and human immunodeficiency virus^19^, which can potentially enhance the disease. Indeed a prior study on SARS-CoV implicated ID sites in antibody-dependent enhancement (ADE)^27,29^. Recent research showed that passive immunisation with early convalescent COVID-19 serum in hamsters resulted in significantly lower viral loads in the respiratory tract without apparent differences in the clinical signs and histopathological changes^30^. Even so, these results provide limited information on ID. We still do not know how to effectively target immunodominance, and we still need to understand the mechanism underlying this phenomenon.

Several candidate vaccines against SARS-CoV-2 including mRNA, inactivated virus, and recombinant adenovirus vaccines have started phase I clinical trials in the US and China^8^. However, there are no reports on the targets of these candidate vaccines, and they might have disease-enhancing effects. Our findings provide evidence for using specific linear antigenic epitopes of S protein instead of the entire S protein for vaccines against SARS-CoV-2 with better safety.

## Acknowledgements

We would like to thank the technicians in the laboratories of JD Huang and KY Yuen for their help in running the project. This work was supported by Health and Medical Research Fund (HKM-15-M09), the Shenzhen Peacock Team Project (KQTD2015033117210153), Shenzhen Science and Technology Innovation Committee Basic Science Research Grant (JCYJ20170413154523577) and China Postdoctoral Science Foundation (2019M663167).

## Author Contributions

BZ Zhang, YF Hu, LL Chen, YG Tong, TCC Yau, KKW To, KY Yuen, and JD Huang designed the study. IFN Hung recruited all the patients. BZ Zhang, YF Hu, KH Chan, and LL Chen performed the experiments. JC Hu, JP Cai, Y Dou, J Deng, HR Gong, C Kuwentrai, WJ Li, XL Wang, H Chu, and CH Su participated in the study. BZ Zhang, YF Hu, LL Chen, YG Tong, TCC Yau, KKW To, KY Yuen, and JD Huang analysed the data. BZ Zhang, YF Hu, and JD Huang wrote the manuscript.

## Competing Interests

The authors declare that they have no competing interests.

## Data and Materials Availability

All data used to draw the conclusions in the paper are presented in the paper and/or the supplementary materials.

## Notes

### Competing Interest Statement

The authors have declared no competing interest.

