## Supplementary Materials for "Mapping the Immunodominance Landscape of SARS-CoV-2 Spike Protein for the Design of Vaccines against COVID-19"

Expression and purification of recombinant protein

The genes encoding the spike RBD (amino acid residues 306 to 543 of the spike protein) and full-length NP of SARS-CoV-2 were codon-optimized with E. coli. Detail information about two recombinant proteins refer to previous work[1].


**Figure S1. Detection of IgG, IgM and IgA-specific antibody titre for NP and SP_RBD in convalescent sera from COVID-19 patients by ELISA.** Heat-inactivated sera were three-fold serially diluted. For patient No. 22, 26, 27, 28, 29, 30 and 31, IgA data was unavailable.

**Figure S2. Detection of antibodies specific for S protein mapping of peptides in early convalescent sera from COVID-19 patients by ELISA.**

**Table S1. List of synthesized peptides.**
