## Supplementary figures and images for "Mapping the Immunodominance Landscape of SARS-CoV-2 Spike Protein for the Design of Vaccines against COVID-19"

### Figure S1

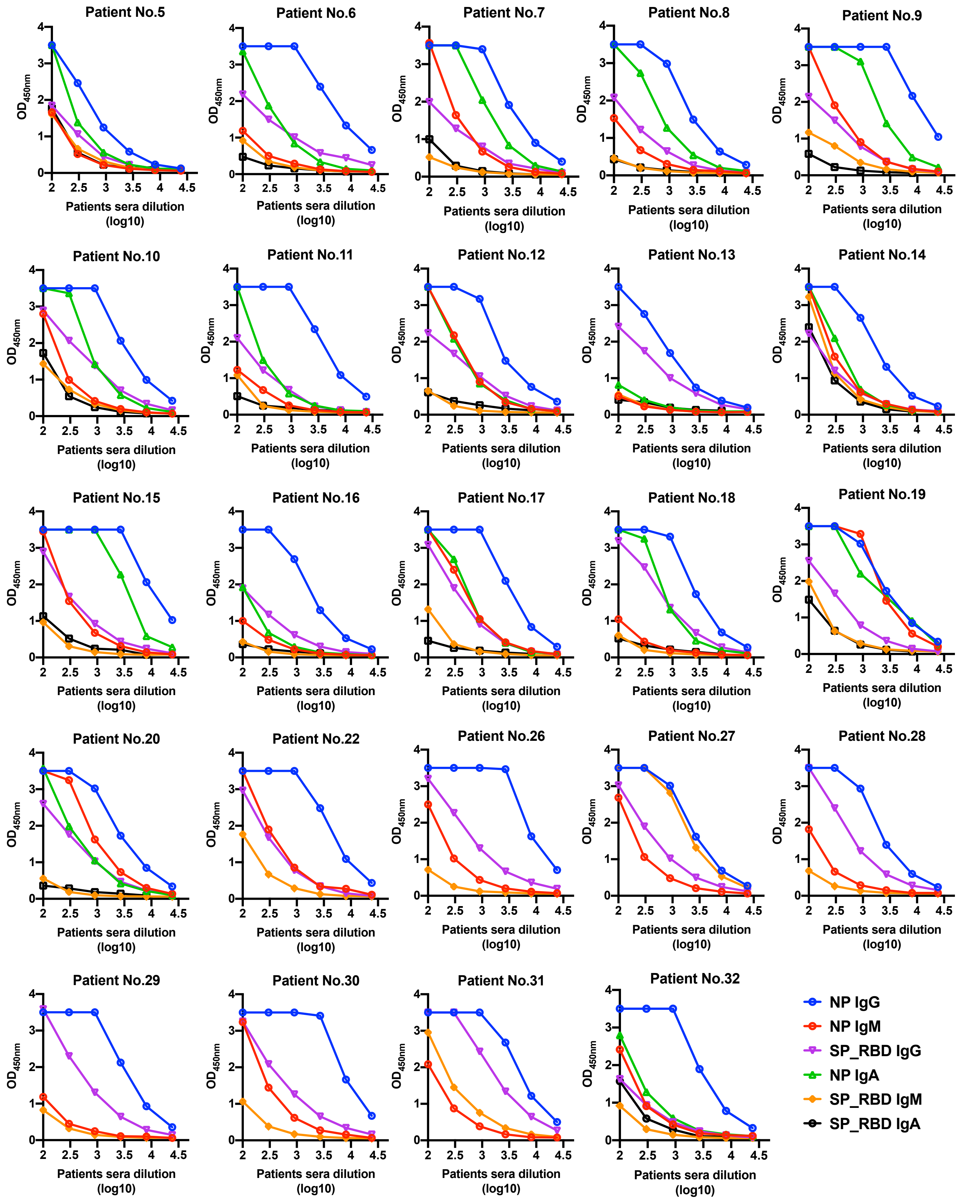

### Figure S2

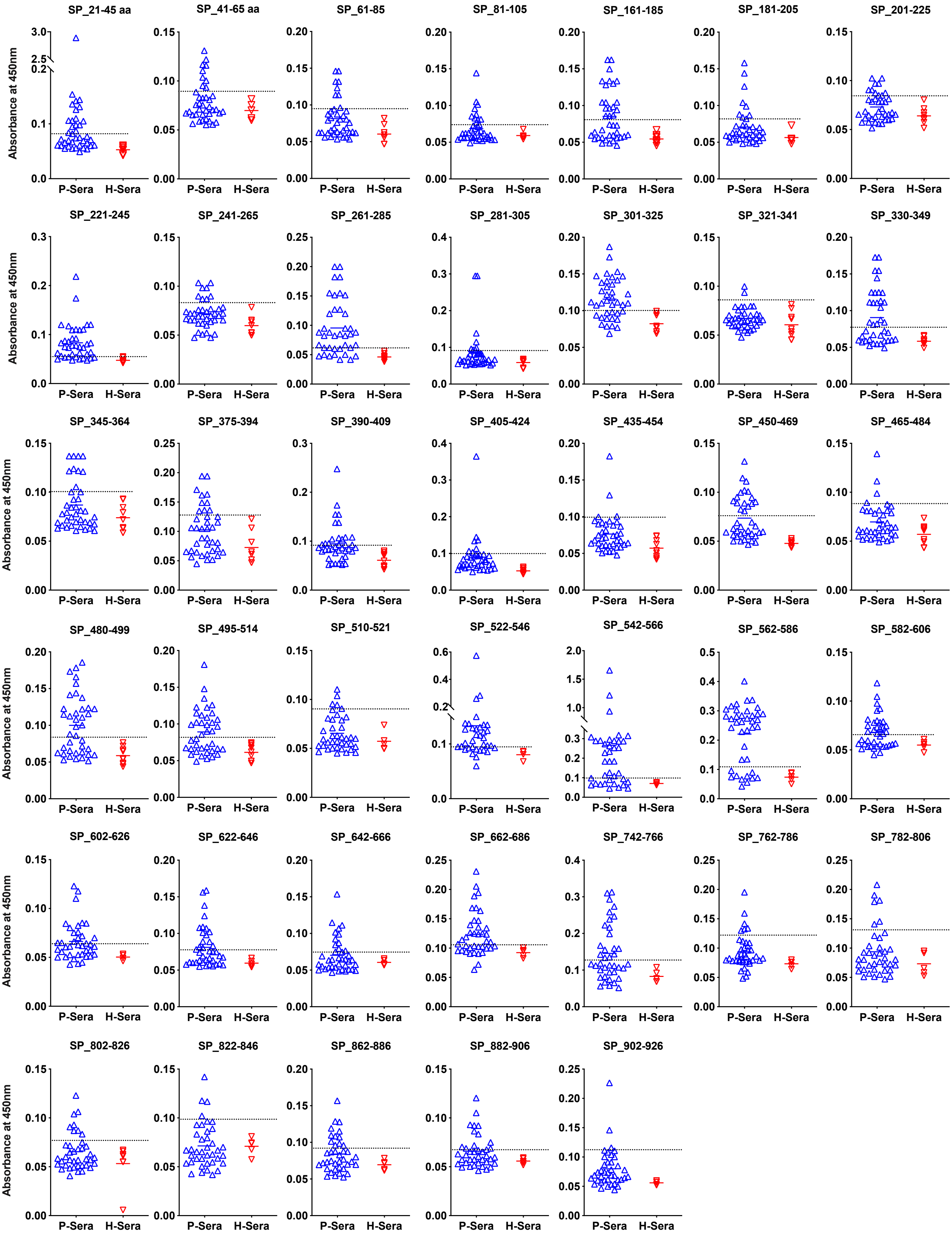
