## Supplementary material for "Mapping the Immunodominance Landscape of SARS-CoV-2 Spike Protein for the Design of Vaccines against COVID-19": Table S1

### Table S1. List of synthesized peptides.

| Position | Sequence |
|---|---|
| 21-45 | RTQLPPAYTNSFTRGVYYPDKVFRS |
| 41-65 | KVFRSSVLHSTQDLFLPFFSNVTWF |
| 61-85 | NVTWFHAIHVSGTNGTKRFDNPVLP |
| 81-105 | NPVLPFNDGVYFASTEKSNIIRGWI |
| 161-185 | SSANNCTFEYVSQPFLMDLEGKQGN |
| 181-205 | GKQGNFKNLREFVFKNIDGYFKIYS |
| 201-225 | FKIYSKHTPINLVRDLPQGFSALEP |
| 221-245 | SALEPLVDLPIGINITRFQTLLALH |
| 241-265 | LLALHRSYLTPGDSSSGWTAGAAAY |
| 261-285 | GAAAYYVGYLQPRTFLLKYNENGTI |
| 281-305 | ENGTITDAVDCALDPLSETKCTLKS |
| 301-325 | CTLKSFTVEKGIYQTSNFRVQPTES |
| 321-341 | QPTESIVRFPNITNLCPFGEV |
| 330-349 | PNITNLCPFGEVFNATRFAS |
| 345-364 | TRFASVYAWNRKRISNCVAD |
| 375-394 | STFKCYGVSPTKLNDLCFTN |
| 390-409 | LCFTNVYADSFVIRGDEVRQ |
| 405-424 | DEVRQIAPGQTGKIADYNYK |
| 420-439 | DYNYKLPDDFTGCVIAWNSN |
| 435-454 | AWNSNNLDSKVGGNYNYLYR |
| 450-469 | NYLYRLFRKSNLKPFERDIS |
| 465-484 | ERDISTEIYQAGSTPCNGVE |
| 480-499 | CNGVEGFNCYFPLQSYGFQP |
| 495-514 | YGFQPTNGVGYQPYRVVVLS |
| 510-521 | VVVLSFELLHAP |
| 522-546 | ATVCGPKKSTNLVKNKCVNFNFNGL |
| 542-566 | NFNGLTGTGVLTESNKKFLPFQQFG |
| 562-586 | FQQFGRDIADTTDAVRDPQTLEILD |
| 582-606 | LEILDITPCSFGGVSVITPGTNTSN |
| 602-626 | TNTSNQVAVLYQDVNCTEVPVAIHA |
| 622-646 | VAIHADQLTPTWRVYSTGSNVFQTR |
| 642-666 | VFQTRAGCLIGAEHVNNSYECDIPI |
| 662-686 | CDIPIGAGICASYQTQTNSPRRARS |
| 742-766 | ICGDSTECSNLLLQYGSFCTQLNRA |
| 762-786 | QLNRALTGIAVEQDKNTQEVFAQVK |
| 782-806 | FAQVKQIYKTPPIKDFGGFNFSQIL |
| 802-826 | FSQILPDPSKPSKRSFIEDLLFNKV |
| 822-846 | LFNKVTLADAGFIKQYGDCLGDIAA |
| 842-866 | GDIAARDLICAQKFNGLTVLPPLLT |
| 862-886 | PPLLTDEMIAQYTSALLAGTITSGW |
| 882-906 | ITSGWTFGAGAALQIPFAMQMAYRF |
| 902-926 | MAYRFNGIGVTQNVLYENQKLIANQ |
